# Full-length structure of the host targeted bacterial effector Bep1 reveals a novel structural domain conserved in FIC effector proteins from *Bartonella*

**DOI:** 10.1101/2024.03.25.586700

**Authors:** Markus Huber, Alexander Wagner, Jens Reiners, Carsten E. Seyfert, Timothy Sharpe, Sander H.J. Smits, Tilman Schirmer, Christoph Dehio

## Abstract

Bacterial effector proteins translocated via a type-IV secretion system (T4SS) typically harbor a C-terminal segment required for recognition by the type-IV secretion coupling protein ^1^. In the α-proteobacterial pathogen *Bartonella*, the signal is bipartite being composed of a BID (Bep intracellular delivery) domain and a positively charged C-terminal tail ^2^. Here, we show the crystal structure of full length *Bartonella* effector protein 1 (Bep1), which shows a novel FIC – OB – BAS(BID) domain arrangement conserved in the majority of Beps with the BID domain inserted into the newly discovered BAS parent domain. We propose that the BAS domain is necessary for the overall “boomerang”-like shape of Bep1 and that it plays a role during translocation through the T4SS.

## INTRODUCTION

The □-proteobacterial genus *Bartonella* comprises arthropod-borne facultative, intracellular pathogens that are adapted to mammals and frequently cause disease (e.g. cat scratch disease, bacillary angiomatosis or peliosis) in humans ^3^. A vast majority of *Bartonella* spp. utilize a VirB/VirD4 Type-IV secretion system (T4SS) to translocate an arsenal of *<u>B</u>artonella* <u>e</u>ffector <u>p</u>rotein<u>s</u> (Beps) into mammalian host cells. Multiple Beps have been characterized and known functions include the inhibition of host cell apoptosis by BepA ^4^, triggering of F-actin driven cytoskeletal processes by BepC ^5^ and the selective targeting of Rac GTPases by Bep1 ^6^.

Beps are multi-domain proteins which share a common architecture at their C-terminus, consisting of a ∼120 amino acid long four-helix bundle, termed <u>B</u>ep intracellular <u>d</u>elivery (BID) domain, and a positively charged tail of variable length ^2, 7^. The two elements constitute a bipartite C-terminal secretion signal that mediates translocation via the VirB/VirD4 T4SS and is evolutionary conserved not only in the Beps of *Bartonella*, but also in the conjugal DNA-transfer-related relaxases and in toxins encoded by many □-proteobacterial species ^2, 8, 9^.

The N-terminal part of Beps is more divergent and can be composed of additional BID domains with secondarily evolved effector functions ^10, 11^, or tyrosine phosphorylation motifs that act as scaffolds to recruit host cell signaling proteins ^12^. However, more than 70% of all Beps possess an N-terminal FIC (<u>F</u>ilamentation induced by <u>c</u>AMP) domain and an OB (oligonucleotide binding) fold (FIC-OB Beps) preceding the BID domain ^13, 14^. Most FIC domains mediate posttranslational modifications such as AMPylation and are folded to a core composed of six □-helices ^15–17^.

*Bartonella* utilize the VirB/VirD4 T4SS ^18^ for the translocation of Beps into eukaryotic host cells ^2, 19, 20^. The T4SS between different organisms are structurally related and are minimally composed of 12 subunits termed VirB2-11 and VirD4 (referring to the nomenclature of the paradigmatic *Agrobacterium tumefaciens* VirB/VirD4 T4SS) ^21^. While VirB2-11 are crucial for the assembly of the VirB/VirD4 T4SS, the type-IV secretion coupling protein (T4CP) VirD4 binds substrates prior to their translocation ^22^. It has also been shown, that binding of substrates to the T4CP can require additional accessory proteins, e.g., in *L. pneumophila* ^23–26^. The translocation route through the T4SS is narrow to an extent that proteinaceous substrates require unfolding for efficient translocation, as has been shown for the 107-kDa relaxase TrwC ^27^, CagA ^28^ and Icm/Dot-translocated substrates ^29^. How substrates are unfolded is yet unknown. Recently, we identified the T4SS effector Bep1 from *Bartonella rochalimae* to selectively target and AMPylate Rac GTPases via its FIC domain ^6^. Bep1 and its many homologues present in pathogenic *Bartonella* spp. were believed to have a canonical FIC-OB-BID architecture. In this study, we describe the full-length structure of Bep1 from *Bartonella clarridgeiae* as the first complete structure of a Bep-T4SS-effector. In the C-terminal part, that had previously been described as unstructured region ^30, 31^, the Bep1 structure revealed a domain with a novel fold into which the BID domain is inserted. Sequence analyses showed that the domain is confined to FIC domain-containing Beps and to □-proteobacterial toxins associated with T4SSs. Due to its apparent function as a scaffold, we termed the new domain BAS (BID Associated Scaffold domain). In addition, we show that Bep1 undergoes temperature-dependent conformational changes, and partially unfolds under physiological temperatures, which might be a prerequisite for effective translocation.

## RESULTS

### Bep1 is monomeric, active and adopts a boomerang-like shape

To gain mechanistic and structural insights into Bep1, we purified full-length Bep1 (558 amino acids, tMw: 63 kDa) from *Bartonella clarridgeiae* to homogeneity (inlet Figure 1A). In size-exclusion chromatography coupled with multi-angle light-scattering (SEC-MALS) experiments, Bep1 eluted as a monomer (Figure 1A, Table S2). In addition, we observed a small shoulder in the elution profile corresponding to a species with the approximate molecular mass of monomeric Bep1_FIC-OB_ (Bep1_1-309_), suggesting proteolytic processing as had been observed for BepA before ^15^ (Figure S1A). *In vitro*, full-length Bep1 AMPylates small GTPase Rac1 (Figure S1B), as has been shown previously for the Bep1_FIC-OB_ fragment ^6^.

**Figure 1:**
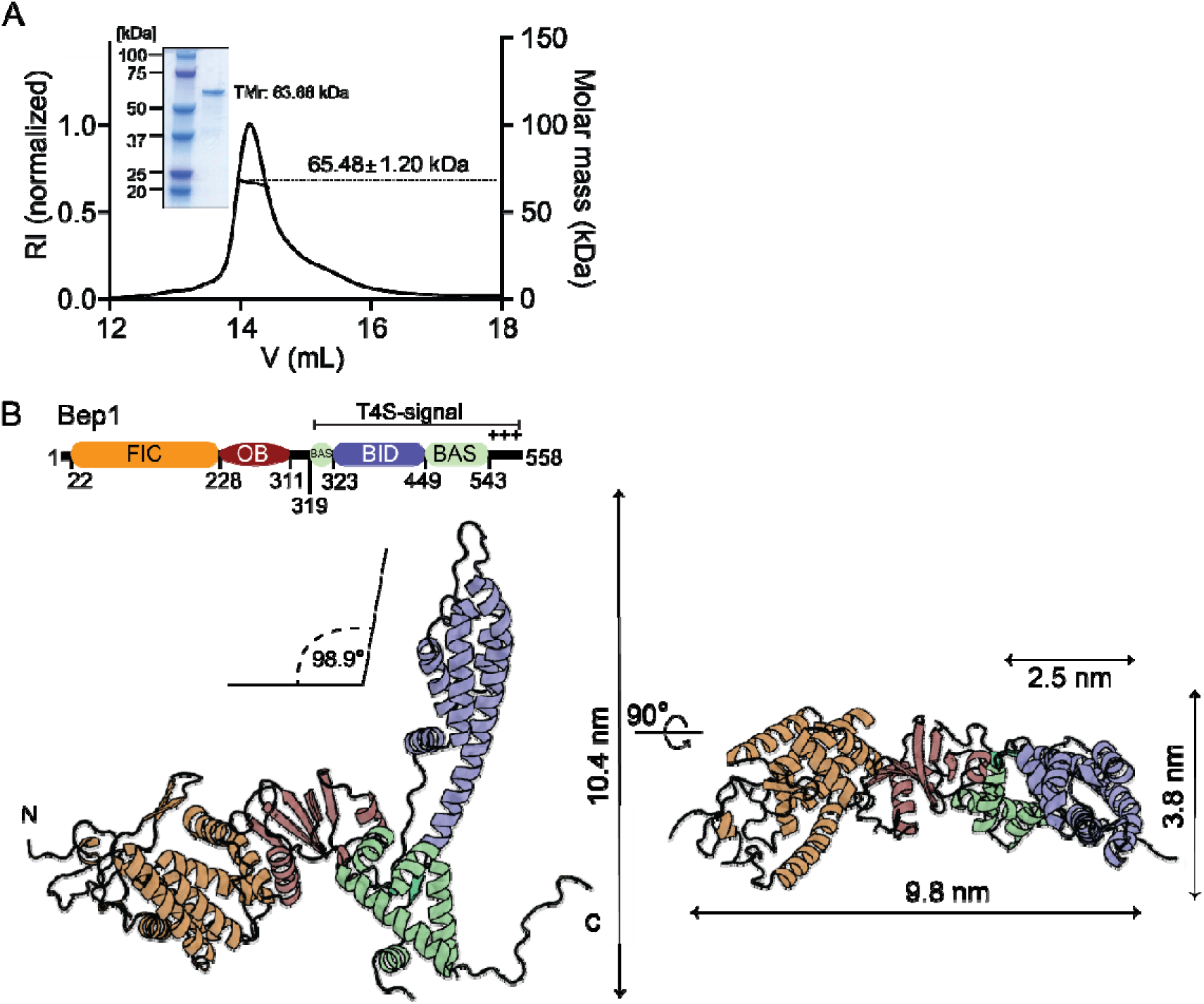
Crystal structure of full-length Bep1. (**A**) Size-exclusion chromatography coupled multi-angle light scattering (SEC-MALS) profile of Bep1 using a GE Healthcare10/300 Superdex 200 increase column. Bep1 elutes with an apparent molecular mass of 65 kDa. The inlet depicts the Coomassie stained SDS-gel of purified Bep1. RI-refractive index. TMr = theoretical Molecular mass. Dots indicate corresponding molar mass (kDa). **(B)** Overall structure of Bep1 (7ZBR) composed of a filamentation induced by <u>c</u>AMP domain (FIC, orange), an <u>o</u>ligonucleotide <u>b</u>inding fold (OB, red) which precedes the <u>B</u>ep intracellular <u>d</u>elivery <u>fold</u> (BID, blue) and the discontinuous <u>B</u>ID <u>a</u>ssociated domain (BAS, lime green).

Bep1 crystallized in space-group P3_2_21 and the structure was solved to a resolution of 3.0 Å by molecular replacement followed by alternating cycles of model building and refinement to a final R_work_ = 27% and R_free_ =30% (see STAR Methods and Table S1 for details). Continuous electron density defines the main-chain from residues 16 to 558, with the exception of residues 470 to 481. The structure adopts an L-or boomerang-shape formed by two wings of roughly 10 nm in length that form an angle of approximately 100° (Figure 1B). The multi-domain structure is composed of FIC, OB and BID folds, with the latter domain found inserted into the BAS domain, which exhibits a novel fold.

The structure of the FIC-OB part of the full-length protein is virtually identical to the previously determined structure of the respective construct of Bep1_FIC-OB_ (PDB 4NPS), with a root-mean-square deviation (RMSD) of 0.56 Å for 241 C□ atoms (Figure S1C). Thus, the fold of the FIC domain and the OB-fold are not altered by the presence of the BAS and BID domains and the remaining part of the peptide chain. The BID domain of Bep1 is similar to the previously solved isolated BID domain structures ^7^. For example, Bep1_BID_ (Bep1_328-447_) aligns with *Bro*Bep6_tBID1_ (PDB 4YK1) with an RMSD of 1.40 Å for 87 Cα atoms, although the sequence identity is very low with 21% (Figure S1D).

### The BAS domain: compact and highly conserved

The BAS domain of Bep1 consists of five anti-parallel α-helices and a two-stranded antiparallel β-sheet. The domain extends from residues 320 to 543, but with residues 324 to 449 belonging to the BID domain, which is found inserted between the two β-strands (Figure 2A). A compact hydrophobic core (Figure 2B) is formed by helices α2 to α5 and the β-sheet. There are no homologous full-length structures of Bep1 known to date and we found no significant structural homologs of the BAS domain in the Protein Data Bank as screened by DALI (Holm 2020) and no significant sequence homologs as scanned by ScanProsite (de Castro, Sigrist et al. 2006).

**Figure 2:**
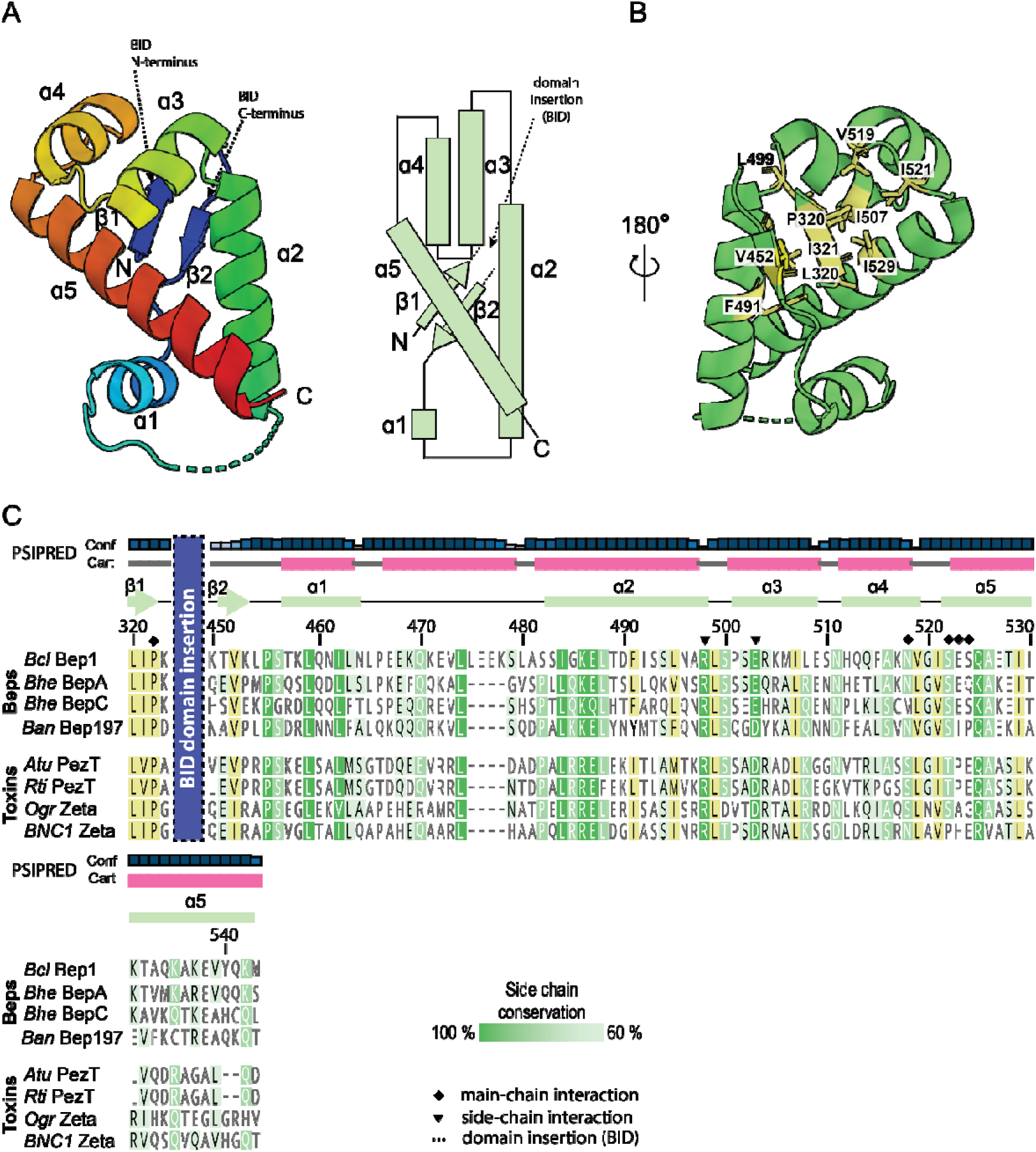
Structure and sequence conservation of the BAS domain. (**A**) Structure (left) and topology (right) of the BID associated (BAS) domain formed by 5 α-helices and a 2-stranded antiparallel β-sheet. In the full-length protein, the BID domain is found inserted between β1 and β2 as indicated. Note, that the stretch be-tween α1 and α2 (470-481) is not resolved in our structure. **(B)** The BAS domain turned by 180° about a vertical axis with respect to the view in panel A, with conserved hydrophobic residues shown as yellow sticks. **(C)** Sequence alignment of the Bep1 BAS domain with other FIC-OB Beps and □-proteobacterial toxins possessing a BID domain. Secondary structure elements of Bep1 BAS are shown on the top (observed in the structure, green; predicted by PSIPRED ^45, 46^, pink). Highly conserved residues (100% identity) are shown in white with lime-green background. Conserved residues (>80% identity) are white with a lighter green background. Partially conserved residues (>60% identity) are black with a light green background. Residues involved in intra– and inter-domain interactions (see Figure 4) are highlighted with a black triangle (side-chain) or diamond (main-chain) on top of the alignment. Conserved hydrophobic residues are shown with yellow background.

Based on sequence comparison, the BAS domain appears to be well conserved among FIC-OB Beps and □-proteobacterial toxins that are associated with the VirB/VirD4 T4SS (Figure 2C). The secondary structure of BAS agrees well with a prediction using PSIPRED (http://bioinf.cs.ucl.ac.uk/psipred/) (Jones 1999, Buchan and Jones 2019) with the unresolved stretch between α1 and α2 (residues 470 to 481) predicted to form an additional helix.

### BID as a BAS insertion domain

According to the classification in Stanger et al. (2017) ^7^ Bep1 is an ancestral tBIDx *Bartonella* effector protein. For this class, sequence conservation has been noted at both BID termini (Figure 3 in there). The full-length Bep1 structure shows that these segments (L320(I/V)P and V452xxP in Bep1) in fact belong to the structural BAS parent domain. They contribute to its hydrophobic core and constitute part of its central β-sheet. Figure S2C shows that such conservation is noticeable also in some α-proteobacterial toxins indicative of the presence of a BAS domain. The structural variations found for BID domains at their chain termini, both amongst the known isolated domain structures ^7^, as well as with respect to the full-length Bep1 structure (Figure 3), can safely be attributed to the missing structural context in the truncated fragments.

**Figure 3:**
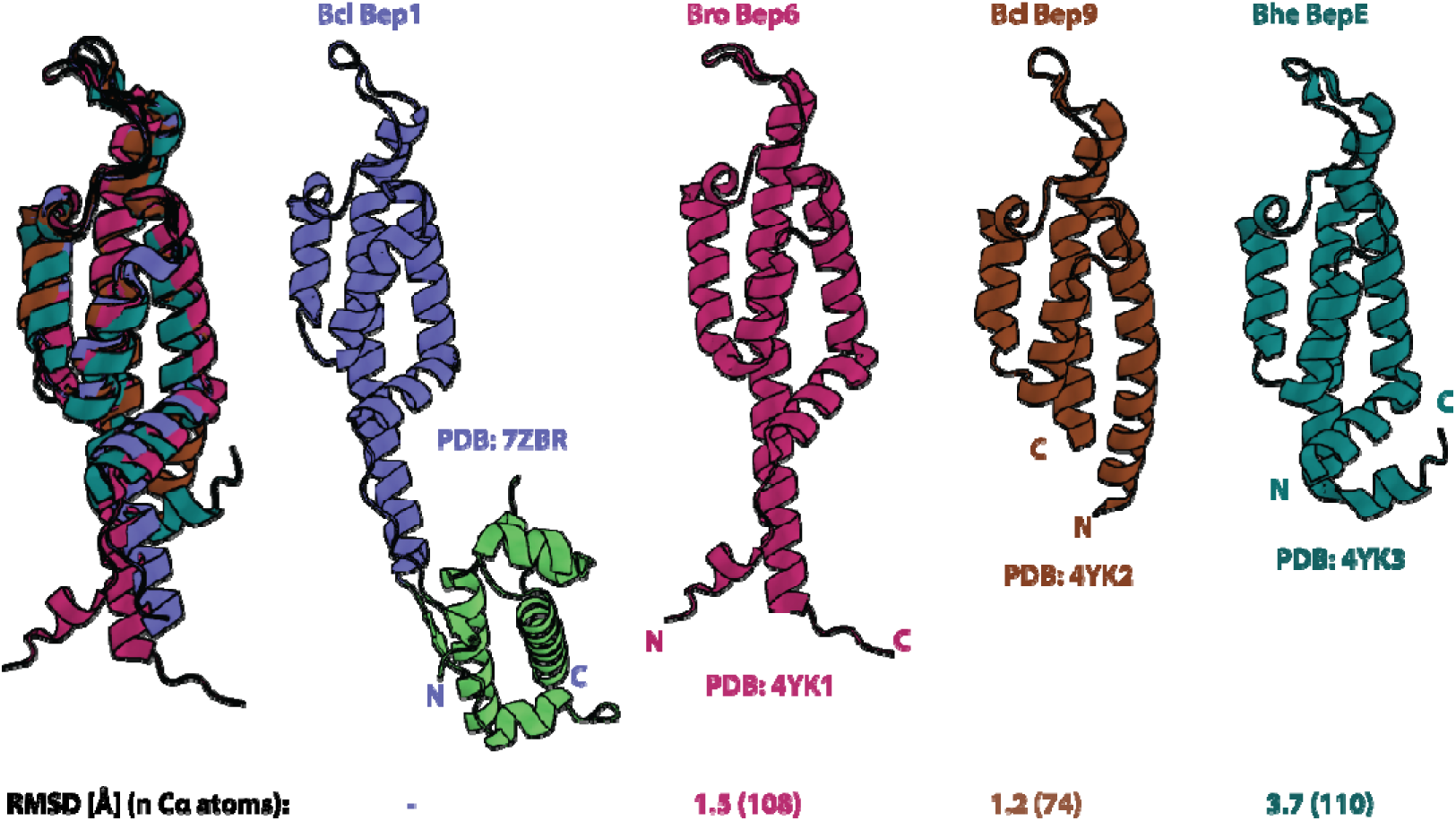
Termini of isolated BID domains show structural variation and are tethered together by the BAS domain in full-length Bep1. Superposition (left) and side-by-side view of BID domain structures. For Bep1, also the BAS domain (green) is shown. The crystal structure of Bep9 from *B. clarridgeiae* (brown) is of isoform 3 (Bep9/3, see Figure S3). RMSD values refer to the comparison with Bep1.

### Interactions between the Bep1 domains

To understand the boomerang-like shape conformation of Bep1, we analyzed the contacts between the different domains (see also Figure 1). While the OB-BAS contact is by far the largest with a buried surface area of about 1300 Å^2^, the BID domain forms a contact of about 650 Å^2^ with its BAS parent domain and a small contact (270 Å^2^) with the OB domain.

Relevant polar interactions are shown in Figure 4A-D. Conspicuously, many of the interactions are ionic and include capping of helices α4 and α5 of the BAS domain by the OB residues Lys298 (Figure 4A) and Asp274 (Figure 4B), respectively. Glu324 of the BID domain plays a central role in that it can form salt-bridges with both Lys298 of the OB domain and Arg498 of the BAS parent domain (Figure 4C and D, respectively). In addition, there is another BID:BAS salt-bridge between Arg447 and Glu503 (Figures 4D). Almost all aforementioned residues are fully conserved (Figures 2C and S2). Taken together, the multiple inter-domain interactions suggest that the BAS domain is responsible for the relative arrangement of the OB and BID domains and, thus, for the overall “boomerang” shape of Bep1.

**Figure 4:**
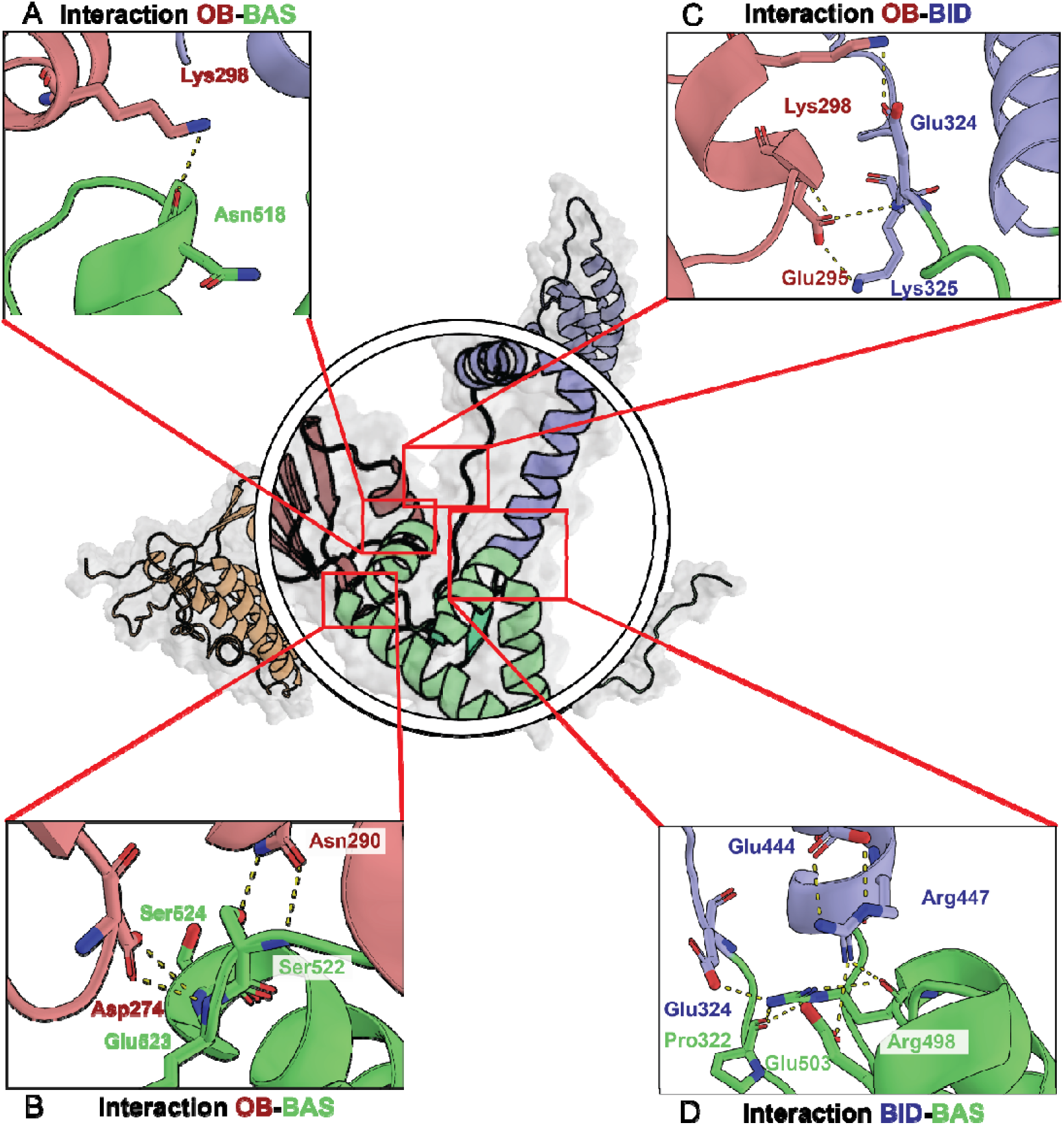
Domain interactions within Bep1. (A-D) Inter-domain interactions of Bep1. Hydrogen bonds and salt-bridges are drawn as yellow dashed lines.

### Destabilization of Bep1 domain arrangement at physiological temperature

To gain further mechanistic and structural insights into T4SS-effectors, like Bep1, we performed small-angle X-ray scattering (SAXS), to identify conformational changes and/or partially unfolding under increasing but still physiological temperatures. SAXS is becoming a common technique to analyze conformational changes upon substrate binding or due to changes in the environment, as well as concentration dependent oligomerization ^32–34^.

We initially performed the SAXS experiment at 15°C, on the Xenocs Xeuss 2.0 with Q-Xoom system. The corresponding SAXS data revealing an *R_g_* and *D_max_*value of 3.95 nm and 13.43 nm, respectively (Figure S5A,C and Table S2). The calculated GASBOR fit showed a χ^2^ value of 1.14, indicating a good agreement with the experimental data (Figure S5A and Table S2). Superimposition of the Bep1 structure and the calculated GASBOR model was done with SUPCOMB and showed a well-fitting overlay of the Bep1 domains FIC (orange), OB (red), BID (blue) and BAS (green) with the SAXS model (Figure 5A). Comparing the theoretical scattering curve of the Bep1 structure via CRYSOL offers a χ^2^ value of 1.38 for the 15°C sample, indicating a good agreement of the structure with the measured scattering data (Figure S5E and Table S2).

**Figure 5:**
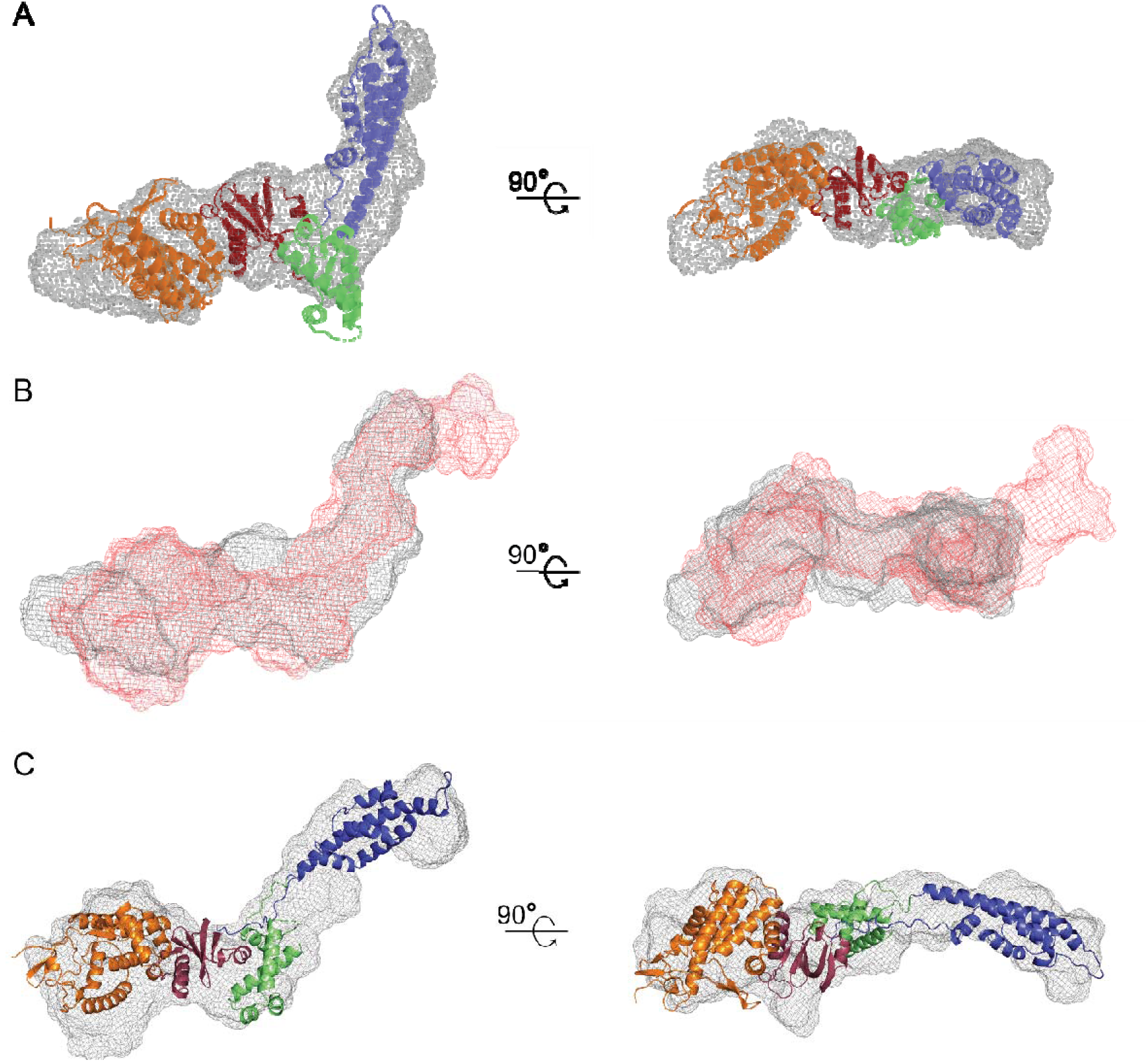
Bep1 GASBOR model. (**A**) The volumetric model from GASBOR is shown as grey mesh, calculated from the Bep1 at 15 °C scattering data. Superimposing of the Bep1 crystal structure was done using SUPCOMB. The Bep1 components FIC domain (orange), OB-fold (red), BID domain (blue) and BAS domain (green) are shown in cartoon representation. **(B)** Overlay of the GASBOR model at 15 °C in grey mesh and from 35 °C in red mesh. **(C)** GASBOR models of 35 °C in grey mesh overlaid with the refined Bep1 model.

To determine conformational changes under elevated temperatures, we heated the sample up elevating the temperature to 35°C to analyze the effects on the Bep1 sample. To avoid prolonged exposure times at high temperature we performed these temperature experiments for Bep1 on the P12 beamline (PETRA III, DESY Hamburg ^35^). The collected SAXS data were analyzed for changes in the particle size (Figure S5B-D and Table S2). By comparison of the analyzed data, we could clearly see that the *R_g_* (3.95 to 4.43 nm) as well as the *D_max_* value (13.43 to 14.42 nm) changes with the temperature rising from 15°C to 35°C. Furthermore, the dimensionless Kratky plot revealed a slightly higher flexibility indicated by the higher *sR_g_* values for the 35°C sample (Figure S5D), but not a complete unfolding of the Bep1 protein. This is in-line with the changes in the particle size, indicating an elongation of the protein (Table S2). We calculated GASBOR models from the different temperatures and compared them to the 15°C model and the crystal structure. The overlay of the 15°C and 35°C GASBOR model, shown in Figure 5B in grey and red mesh representation, visualizes the elongation of the Bep1 protein at higher temperature. Going even higher than 35°C, leads to a rapid aggregation of the protein. Comparing the theoretical scattering curve of the Bep1 structure via CRYSOL offers a χ^2^ value of 2.20 for the 35°C sample. (Figure S5E and Table S2). The corresponding residual plot shows that the higher χ^2^ value mainly comes from the mismatch of the low s region. This indicates a rearrangement of the domains, in-line, with the higher *D_max_* values (Figure S5E and Table S2). Taken together this aligns with the hypothesis that Bep1 stretches at elevated temperatures. With SREFLEX we tried an initial normal mode analysis to refine the Bep1 structure for a better agreement with the scattering data at 35°C and fine tune it manually later on (Figure 5D, Figure S5G and Table S2). The refinement offers a small twist of the FIC domain (orange), the OB fold (red), the BAS domain (lime green) and a more stretched BID domain (blue) (Figure 5C) suggesting that some of the interactions between OB-fold, BID domain and BAS do-main (Figure 4) get lost.

## DISCUSSION

Bacteria have evolved a plethora of secretion systems that are critical during pathogenesis or interbacterial killing. These systems secrete different substrates including DNA, peptidoglycan and proteins. Proteins are translocated in a folded conformation by T2SSs ^36^ or at least partially unfolded by e.g. T1SSs ^37^, T3SSs ^38^ and T4SSs ^27^. In this study we report the first structure of a full-length Bep-T4SS-effector, Bep1, showing a boomerang-like shape, with each wing being around 10 nm in length (Figure 1B). Although the *Bartonella* VirB/VirD4 T4SS machinery has not been microscopically visualized, its translocation channel is probably in the range of around 2 nm to 6 nm in diameter analog to the T4SS machinery encoded by plasmid pKM101 that is structurally well characterized^39^. Considering these shapes and diameters, we hypothesize that secretion of Bep1 and homologs by the T4SS requires partial unfolding in line with other proteinous T4SS substrates ^28, 29^.

The factor(s) that contribute to T4SS-substrate unfolding are not known. In T3SS, a dedicated hexameric ATPase recognizes and unfolds substrates in an ATP-dependent manner ^40^. T4-secretion is energized by three ATPases: VirB4, VirB11 and the T4CP VirD4. These three ATPases fulfill diverse functions during translocation, including T4SS-pilus assembly, substrate-secretion (VirB4, VirB11) and – recognition (T4CP) ^18^. Either of these ATPases could function as an unfoldase for T4SS-substrates and induce the required conformational changes in Bep1. The boomerang shape of Bep1 is held together by hydrophobic interactions and a complex network of conserved ionic bonds. However, interdomain interactions are most numerous between BAS and its inserted BID domain, and between BAS and the OB-fold (Figures 2B and 4), suggesting a scaffolding role for the BAS domain. Binding of Bep1 to either of the aforementioned ATPases and/or other T4SS components might introduce slight rearrangements that could ultimately promote extensive conformational changes or partial unfolding along the scaffold between the BID domain and the OB-fold.

This is supported by the intrinsic flexibility of the effector, that translates through the “knee” shaped by the BAS domain. In our SAXS-experiments we observed that with increasing temperature the angle between the wings of the Bep1 boomerang became wider (Figure 5C), indicating that interactions of the OB-fold and BID domain with the BAS domain are less stable at the physiological temperatures of *Bartonella* host organisms. Overall, Bep1 adopts a more stretched conformation at 35°C compared to 15°C (Figure 5B,C). As 35°C is closer to the mammalian body temperature, the stretched Bep1 conformation could biologically be more relevant with respect to T4-secretion through the bacterial membrane. Our SAXS data furthermore show that Bep1 is partially unfolded at 35°C without the action of an external factor, which might provide evidence of the involvement of intrinsic characteristics of T4SS-effectors in T4-secretion (Figure S5C). External factors, for example the binding of parts of the T4SS to exposed parts of the BAS domain might contribute further to the elongation of Bep1. *In vitro* translocation experiments comparing Bep1 secretion with more stable Bep1 derivatives in combination with SAXS measurements could proof the role of the BAS domain in T4-secretion.

The C-terminal part of Bep1 following the BID domain had previously been described as unstructured region with a positively charged tail required for efficient translocation ^2, 8^. Here we have shown that the major part of this tail together with the short segment housing the L_320_(I/V)P_322_-motif, which precedes the inserted BID domain, forms the well-structured BAS domain. Thereby, the short segment forms one of the β-strands of the β-sheet complementing the hydrophobic core of the BAS domain.

Insertions of a child domain into a parent domain have first been identified in the early 90s ^41^ and have since been found in 9% of multi-domain proteins of the nonredundant Protein Data Bank ^42^.

We speculate that during evolution the BID child domain has been inserted into a loop between β1 and β2 of the BAS parent domain. The conservation of the L_320_(I/V)P_322_-motif in sequences containing the BAS domain, but not in BID domains without a BAS domain (e.g. Bep9 and BepE; Figures 3 and S3) is a strong indication for this insertion event. While the BAS domain might play a role for secretion or as part of the bipartite secretion signal ^2^, the occurrence of non-terminal BID:BAS domain combinations (eg. Bep197 BID1) suggests a function besides that. It is conceivable, that phenotypes attributed to the BID domain could be triggered in association with the BAS domain, as most BID domain constructs used in previous studies also contained a BAS domain ^10, 43, 44^. Moreover, the role of the BAS domain might be linked to an enzymatic function that has yet to be explored. Future structure-function studies might unravel the role of the BAS domain after translocation into host cells through the T4SS.

## Supplementary

**Supplementary Figure 1:**
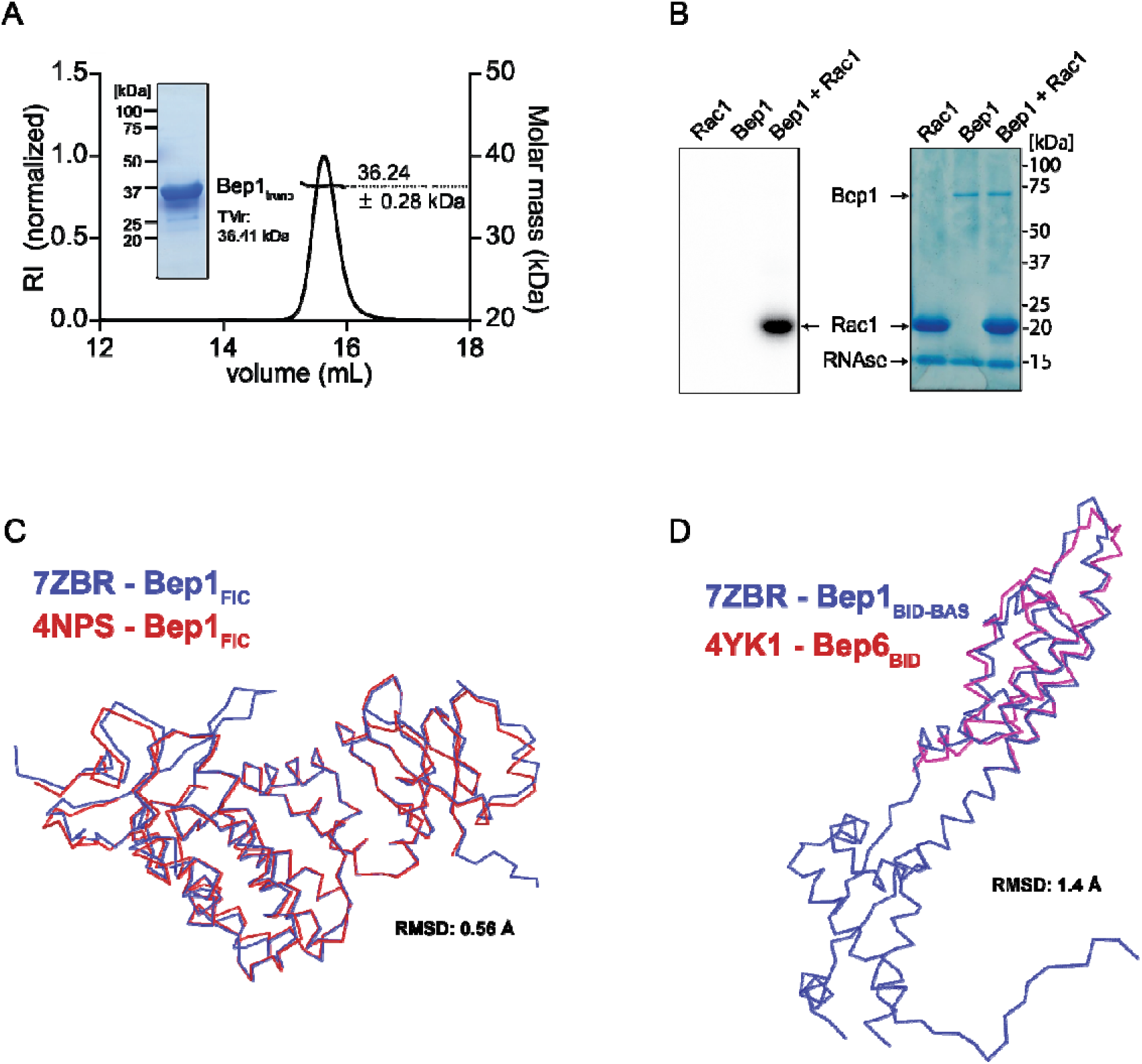
Characteristics of full-length Bep1 (A) Size-exclusion chromatography coupled multi-angle light scattering (SEC-MALS) profile of Bep1_trunc_ using a GE Healthcare10/300 Superdex 200 increase column. Bep1_trunc_ elutes with an apparent molecular mass of 36 kDa. The inlet depicts the Coomassie stained SDS-gel of purified Bep1_trunc_. RI = refractive index. TMr = theoretical Molecular mass. Dots indicate corresponding molar mass (kDa). **(B)** Autoradiogram and SDS-gel of Bep1 and Rac1 after incubation with ^32^P-α-ATP. **(C)** Cα-trace of Bep1FIC-OB from full-length *B. clarridgeiae* (blue) overlayed onto Bep1FIC-OB from *B. rochalimae* (red). **(D)** Cα-trace of Bep1BID-BAS from full-length *B. clarridgeiae* (blue) overlayed onto Bep6BID from *B. rochalimae* (red)

**Supplementary Figure 2:**
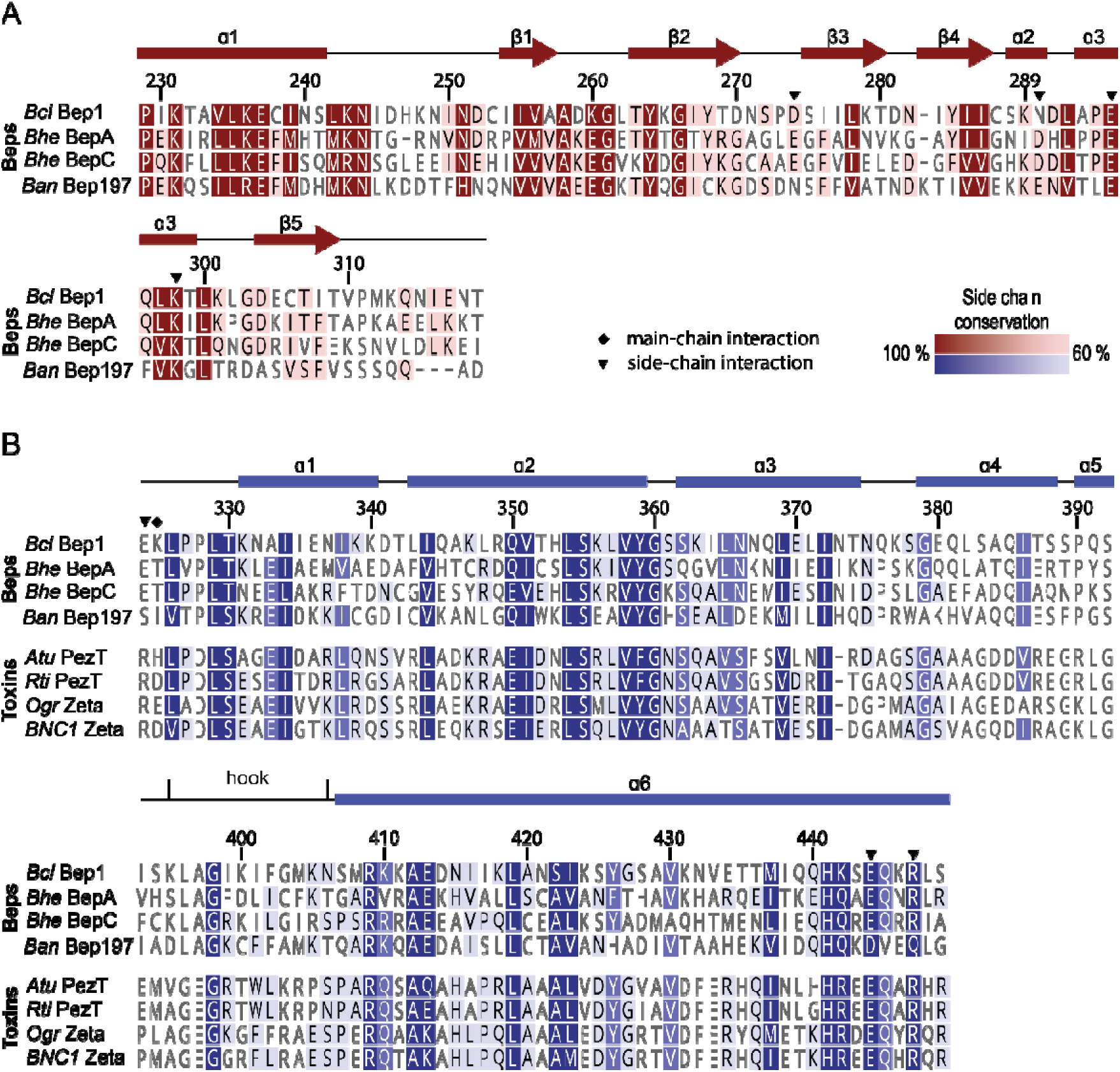
OB-fold and BID domain sequence alignments (A, B) Sequence alignment of the Bep1 OB-fold **(A)** with other Beps and of the Bep1 BID domain **(B)** with other Beps and □-proteobacterial toxins possessing a BID domain. The secondary structure, as observed in the structure, is drawn on top of the alignments. The conservation level (100% – 60%) is indicated by the strength of the respective colour. Residues involved in inter-domain interactions (see Figure 4) are highlighted with a black triangle (side-chain) or diamond (main-chain) on top of the alignment.

**Supplementary Figure 3:**
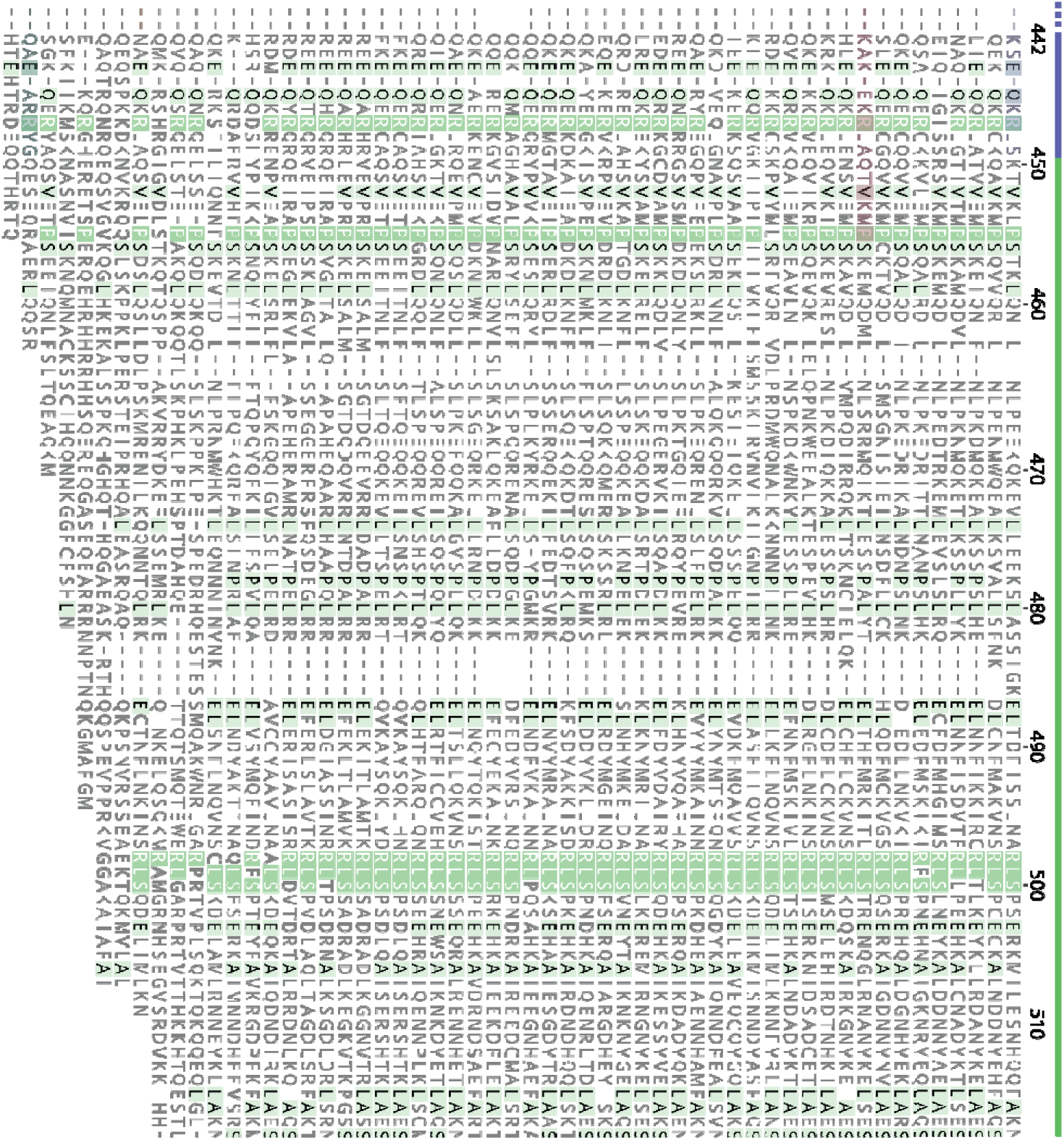
BAS domain sequence alignment of BID domain proteins. Multiple-sequence alignment of BID domain containing Beps and □-proteobacterial toxins zoomed in on the BAS domain and the flanking ends of the respective BID domains. Sequence coverage of structures seen in Figure 3 is indicated with semi-transparent bars of the respective colour. The conservation level (100% – 60%) is shown by the strength of the green background colour.

**Supplementary Figure 4:**
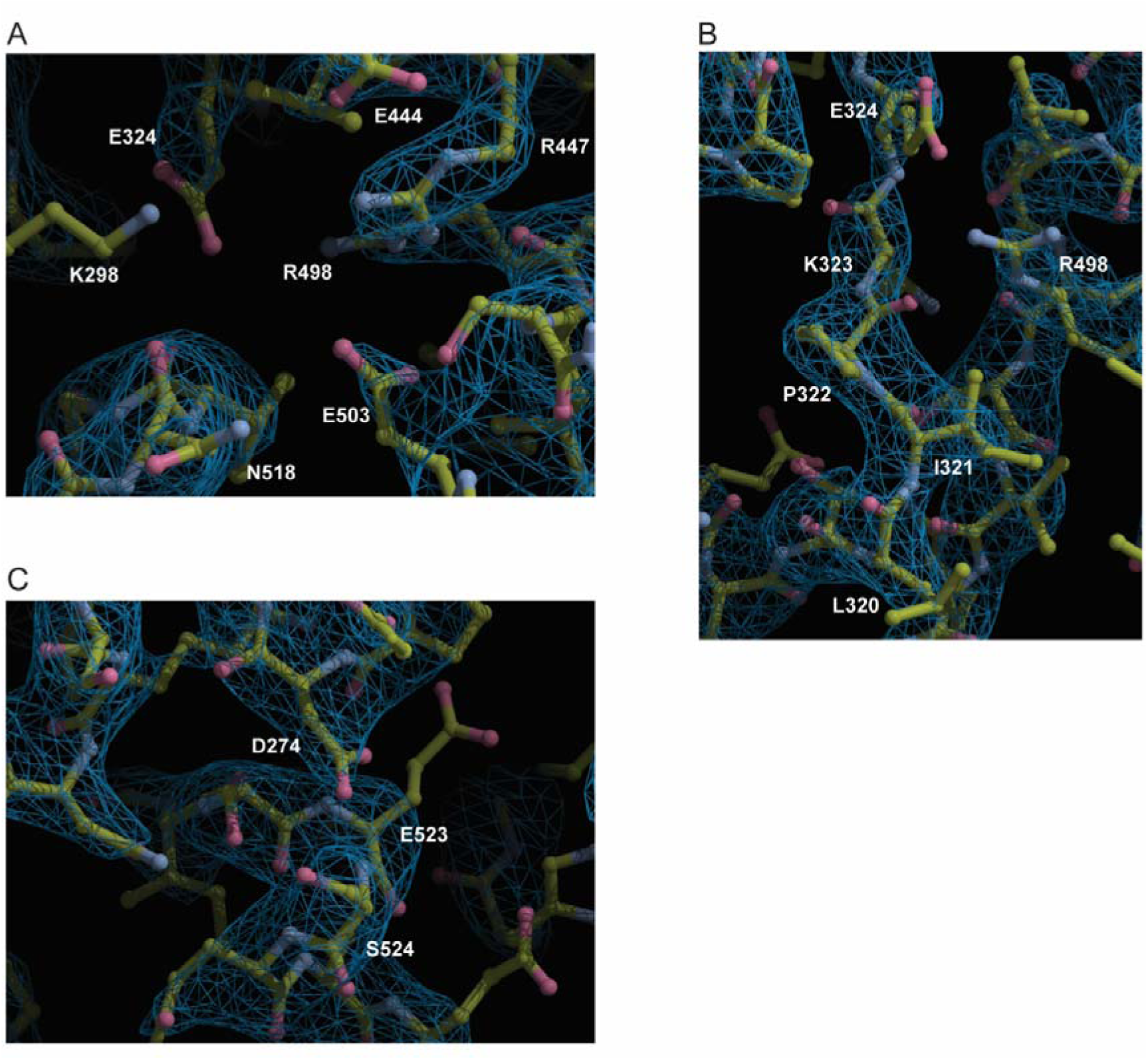
Electron density maps of intra– and interdomain interactions. A-C show electron density with corresponding model of areas from the OB-fold, BAS domain and BID domain involved in interactions (see Figure 4). Maps are shown as meshes with a contour level of 1.9 e/Å^3^ in coot ^47^

**Figure S5:**
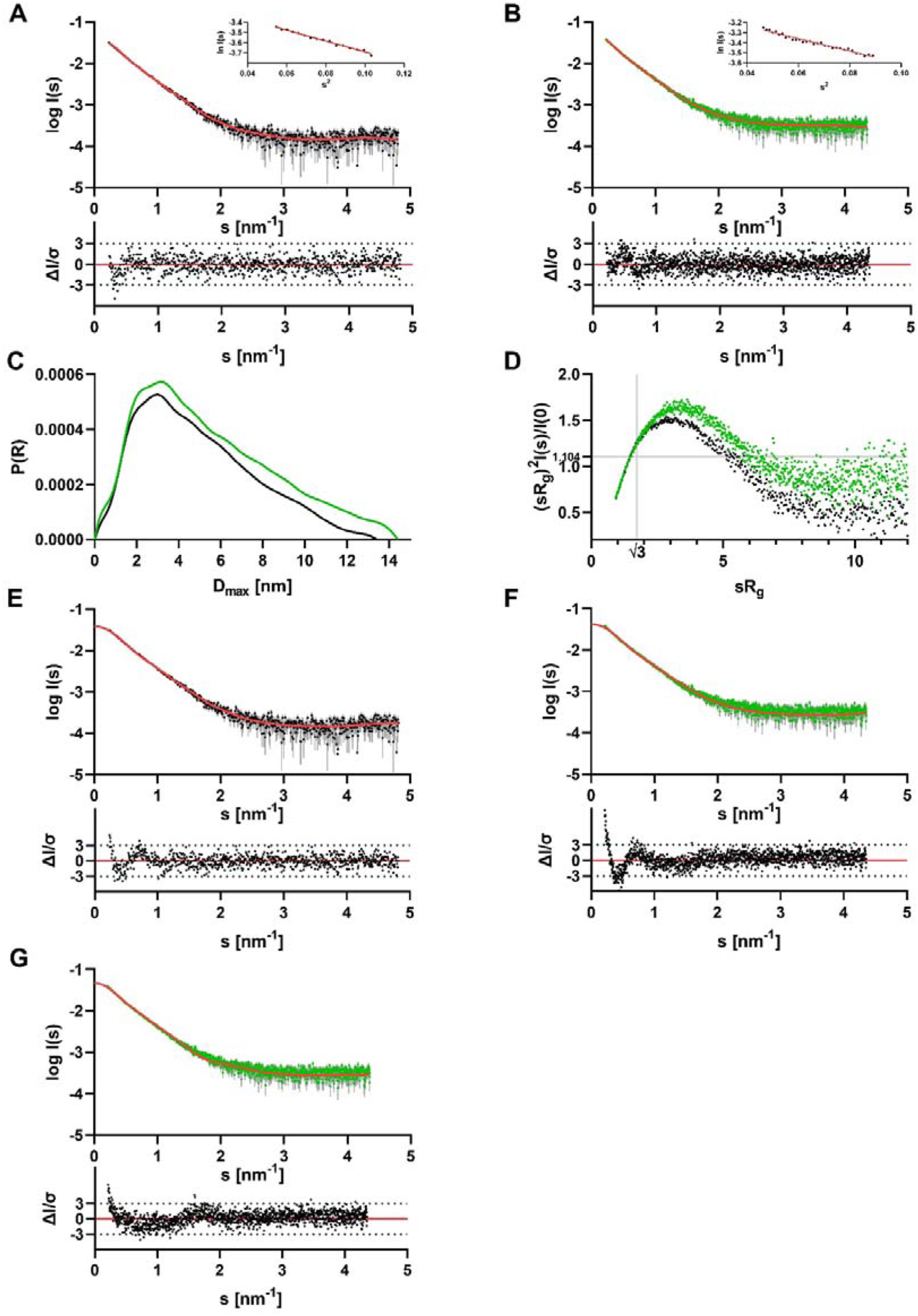
SAXS results for Bep1 at different temperatures. (A) Scattering data of Bep1 at 15°C. Experimental data are shown in black dots, with grey error bars. The ab-initio GASBOR model fit is shown as red line and below is the residual plot of the data. The Guinier plot is added in the right corner. **(B)** Scattering data of Bep1 at 35°C. Experimental data are shown in green dots, with grey error bars. The ab-initio GASBOR model fit is shown as red line and below is the residual plot of the data. The Guinier plot is added in the right corner. **(C)** *p(r)* function of Bep1 at 15°C (black line) and 35°C (green line). **(D)** Dimensionless Kratky plots of Bep1 at 15°C (black dots) and 35°C (green dots). **(E)** CRYSOL fit of the Bep1 crystal structure against the scattering data of Bep1 at 15°C. Experimental data are shown in black dots, with grey error bars. The CRYSOL fit is shown as red line and below is the residual plot of the data. **(F)** CRYSOL fit of the Bep1 crystal structure against the scattering data of Bep1 at 35°C. Experimental data are shown in green dots, with grey error bars. The CRYSOL fit is shown as red line and below is the residual plot of the data. **(G)** CRYSOL fit of the refinement of the Bep1 crystal structure against the scattering data of Bep1 at 35°C. Experimental data are shown in green dots, with grey error bars. The CRYSOL fit is shown as red line and below is the residual plot of the data.

**Table S1:** Data collection and refinement statistics.

|  | Bcl. Bep1 (7ZBR) |
| --- | --- |
| Resolution range | 46.38 - 3.0 (3.107 - 3.0) |
| Space group | P 32 2 1 |
| Unit cell | 70.492 70.492 213.961 90 90 120 |
| Total reflections | 166906 (16534) |
| Unique reflections | 13003 (1271) |
| Multiplicity | 12.8 (13.0) |
| Completeness (%) | 99.63 (98.74) |
| Mean I/sigma(I) | 8.10 (0.66) |
| Wilson B-factor | 110.18 |
| R-merge | 0.1851 (3.604) |
| R-meas | 0.1933 (3.748) |
| R-pim | 0.05439 (1.014) |
| CC1/2 | 0.993 (0.335) |
| CC* | 0.998 (0.709) |
| Reflections used in refinement | 13003 (1255) |
| Reflections used for R-free | 1302 (126) |
| R-work | 0.2699 (0.3957) |
| R-free | 0.3019 (0.4233) |
| CC(work) | 0.924 (0.428) |
| CC(free) | 0.907 (0.238) |
| Number of non-hydrogen atoms | 4196 |
| macromolecules | 4196 |
| ligands | 0 |
| solvent | 0 |
| Protein residues | 531 |
| RMS(bonds) | 0.003 |
| RMS(angles) | 0.62 |
| Ramachandran favored (%) | 98.29 |
| Ramachandran allowed (%) | 1.71 |
| Ramachandran outliers (%) | 0 |
| Rotamer outliers (%) | 0 |
| Clashscore | 2.35 |
| Average B-factor | 102.64 |
| macromolecules | 102.64 |

**Table S2:**
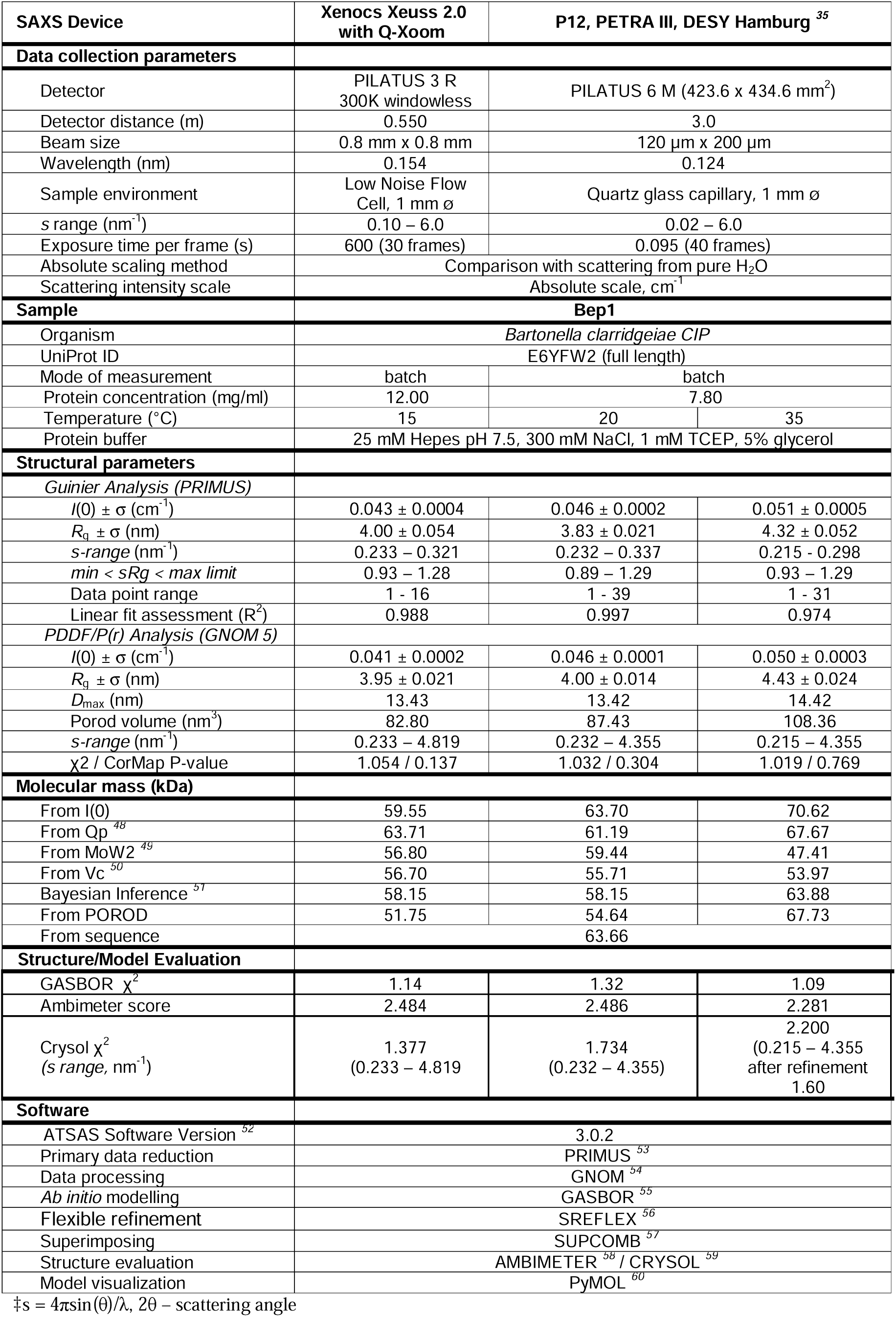
Overall SAXS Data.

## STAR METHODS

### RESOURCE AVAILABILITY

#### Lead Contact

Further information and requests for resources and reagents should be directed to and will be fulfilled by the Lead Contact, Dr. Christoph Dehio.

#### Material Availability

Plasmids generated in this study are available upon request to the Lead Contact.

#### Data and Code Availability

Structural data are deposited on https://www.rcsb.org/ and https://www.sasbdb.org/.

## EXPERIMENTAL MODEL AND SUBJECT DETAILS

### Microbes

*E. coli* cells were cultured at 20°C – 37°C in LB medium supplemented with appropriate antibiotics (Key resource table).

## METHOD DETAILS

### Cloning

A Construct encoding soluble *Bartonella clarridgeiae* (*Bcl*)-Bep1 (Bep1-Fic-OB)_aa1-309_ was cloned via restriction cloning into pRSFDuet™-1, resulting in pAW041. Note that (Bcl)-Bep1 full length _aa1-558_ was received from the Seattle Structural Genomics Center for Infectious Disease (see Key resource table).

### Protein Expression and Purification

Expression plasmids (pCES001, pAW041) were transformed into *E. coli* BL21(DE3). Single colonies were picked to inoculate precultures, respectively, which were incubated overnight at 37°C in 50 ml LB + 1% glucose + 100 μg/ml ampicillin (BG1861-*His6-bep1_bcl_*) or 50 μg/ml kanamycin. Next day, 15 ml of precultures were used to inoculate 1.5 l LB + 1% glucose + the appropriate antibiotic and cultures were grown at 37°C to OD=1.0. Protein expression was induced with 0.2-0.5 mM IPTG and expression cultures were incubated at 21°C for 16 hours. Expression cultures were pelleted, frozen in liquid nitrogen and stored at –80°C. Cell pellets were resuspended in low imidazole buffer (25 mM Hepes pH 7.5, 20 mM imidazole, 300 mM NaCl, 1 mM TCEP, 5% glycerol) supplemented with Benzonase® (Merck) and cOmplete™ Mini EDTA-free protease inhibitor (Roche). Following incubation for 30 min on ice, cells were broken using a French Press (16 000 psi) and supernatant was obtained by centrifugation at 100.000 x g (45 min, 4°C). Subsequently, the supernatant was loaded onto a pre-equilibrated (with low imidazole buffer) HisTrap™excel column (GE Healthcare). Following a wash step of the column with 5 column volumes low imidazole buffer, proteins were eluted with an imidazole gradient from 20 to 500 mM imidazole. Elution fractions were concentrated in Amicon® Ultra-15 centrifugal filters (30 kDa cut-off for Bep1 and sVirD4, 10 kDa cut-off for Bep1-Fic-OB). Concentrated proteins were further purified by Size-exclusion chromatography (SEC) either using a HiLoad 16/60 Superdex 200 pg column (GE Healthcare) or a HiLoad 16/60 Superdex 75 pg column (GE Healthcare) pre-equilibrated in SEC-buffer (25 mM Hepes pH 7.5, 300 mM NaCl, 1 mM TCEP, 5% glycerol). Eluted proteins were frozen in liquid nitrogen and stored at –80°C. Protein concentrations were determined using the Pierce™ BCA Protein Assay kit (Thermo Fisher Scientific) and via direct A280-measurments using a NanoDrop One^C^ UV-Vis spectrophotometer (Thermo Fisher Scientific).

### Multiangle Light Scattering

Size-exclusion chromatography coupled multiangle light scattering (SEC-MALS) of Bep1-derivatives was performed on a GE Healthcare10/300 Superdex 200 increase column, equilibrated overnight with SEC buffer (25 mM Hepes pH 7.5, 300 mM NaCl, 1 mM TCEP, 5% glycerol) at 25°C, using an Agilent 1260 HPLC. 100 ul sample at a concentration of 0.3 mg/ml (Bep1) or 0.2 mg/ml (Bep1-Fic-OB) were applied and elution was monitored an Agilent multi-wavelength absorbance detector (280 nm), a Wyatt Heleos II 8+ multiangle light scattering detector and a Wyatt Optilab rEX differential refractive index detector. 2 mg/ml BSA solution (Thermo Pierce) was injected to calibrate interdetector delay volumes, band broadening corrections, and light scattering detector normalization using the Wyatt ASTRA 6 software (Wyatt Technology). Weight-averaged molar mass was calculated from the light scattering and the differential refractive index (RI) signals using Wyatt ASTRA 6 software (Wyatt Technology).

### Radioactive AMPylation Assay

The AMPylation activity of full-length Bep1 on Rho GTPase, Rac1, was investigated using an *in vitro* AMPylation assay with radioactive phosphate ([α-32 P]-ATP (Hartmann Analytic)) comparable to previously described assay with purified Bep1 FIC domain and bacterial lysates expressing full-length Bep1 ^6^. 50 µM Rho-GTPase (preloaded with GDP), 1 µM purified full-length Bep1 and 1 mM ATP were incubated with 10 µCi [α-32 P]-ATP in reaction buffer (50 mM Tris-HCl, pH 8.0, 150 mM NaCl, 5 mM MgCl_2_, 0.15 mg/mL RNaseA) for 1 h at 30 °C. The reaction was stopped by addition of SDS-sample buffer and heating to 95 °C for 10 min. Samples were separated by SDS-PAGE and subjected to autoradiography.

### Structure determination

For crystallization 12 mg/ml of the purified Bep1 (0.2 mM, supplemented with 5 mM ATP and 5 mM MgCl_2_) were mixed with reservoir solution in a ratio of 1:2 yielding an end concentration of 4 mg/ml. Crystallization was done using the sitting-drop vapour diffusion method by dispensing 0.6 µl in MRC 96-well plates (SWISSCI). The reservoir solution was composed of 100 mM Hepes pH 7.8, 0.175 mM LiCl and 20% v/v PEG 8000. Crystals were obtained at 20°C and frozen in liquid nitrogen with glycerol as an additional cryoprotectant. Data collection was done at the Swiss Light Source of the PSI (https://www.psi.ch/en/sls) on beam-line X06SA (PXI) at λ = 1.0 Å with an EIGER 16M X detector (133 Hz). Images were processed with XDS ^61^. The structure was solved by molecular replacement with Phaser ^62^. As search models the crystal structures of Bep1(FIC-OB) (PDB: 4NPS) from *Bartonella clarridgeiae* and Bep6(BID (PDB: 4YK1) from *Bartonella rochalimae* were used. Model building was done in COOT ^47^ with alternating cycles of refinement in Phenix ^63^. Data collection and refinement statistics are summarized in Table S1.

### Small-Angle X-ray Scattering (SAXS)

We collected the initial SAXS data from Bep1 on our Xeuss 2.0 Q-Xoom system from Xenocs, equipped with a PILATUS 3 R 300K detector (Dectris) and a GENIX 3D CU Ultra Low Divergence x-ray beam delivery system. The chosen sample to detector distance for the experiment was 0.55 m, results in an achievable q-range of 0.10 – 6 nm^-1^. The measurement was performed at 15 °C with a protein concentration range of 3 – 12 mg/ml. The Bep1 sample was injected in the Low Noise Flow Cell (Xenocs) via autosampler. We collect 30 frames with an exposer time of ten minutes/frame and scaled the data to absolute intensity against water. We checked each frame for radiation damage using CorMap/ χ^2^ test, implemented in in PRIMUS ^53^. After checking the frames of the different concentrations we saw no concentration effect and continue the evaluation with the 12 mg/ml data set.

To avoid longer exposer times on high temperature we performed the temperature experiments for Bep1 on the P12 beamline (PETRA III, DESY Hamburg ^35^). The autosampler at P12 was set to the chosen temperature and the Bep1 sample (7.8 mg/ml) was incubated 20 min before measuring. We collected 40 frames for each temperature with an exposer time of 0.095 sec/frame, radiation damage was checked via the SASFLOW pipeline^64^ and identical frames were merged. Data were scaled to absolute intensity against water.

All used programs for data processing were part of the ATSAS Software package (Version 3.0.2) ^52^. Primary data reduction was performed with the program PRIMUS^53^. With the Guinier approximation ^65^, we determine the forward scattering *I(0)* and the radius of gyration (*R_g_*). The program GNOM ^54^ was used to estimate the maximum particle dimension (*D_max_*) with the pair-distribution function *p(r)*. Cross evaluations of the *D_max_* values was also done using SHANUM ^66^. Low resolution *ab initio* models were calculated with GASBOR ^55^. We refined the Bep1 crystal structure with SREFLEX ^56^ using the 35 °C scattering data and fine tune the structure manually later on. Superimposing of the Bep1 structure was done with the program SUPCOMB^57^.The theoretical scattering of the Bep1 structure was computed with CRYSOL ^59^ (ns 501, lm 70, fb 18), using the same s-range like GNOM and compared against the solution scattering data of the different used temperature.

## QUANTIFICATION AND STATISTICAL ANALYSIS

Statistical parameters are indicated in figures and respective legends. Error bars in the SAXS scattering plots (Figure S5) show the standard deviation. Statistical data of the Bep1 structure data collection and refinement are summarized in Table S1. Statistics of the SAXS Data collection are shown in Table S2.

## DATA AND SOFTWARE AVAILABILITY

We uploaded the SAXS data to the Small Angle Scattering Biological Data Bank (SASBDB) ^67, 68^. Protein structure data have been deposited in the Protein Data Bank under accession number 7ZBR. Molecular replacement was done with Phaser ^62^. Several rounds of iterative model building and refinement were performed using Coot ^47^ and Phenix.refine ^63^, respectively. MSA were done with the GENEIOUS software package Version 7.1.7 and later ^69^. Visualization of structures and models were done with pymol ^70^.

## Acknowledgements

We acknowledge DESY (Hamburg, Germany), a member of the Helmholtz Association HGF, for the provision of experimental facilities. Parts of this research were carried out at PETRA III, and we would like to thank Tobias Gräwert (EMBL Hamburg) for the assistance in using beamline P12.

## Funding

The work was funded by project grant 310030B_201273 from the Swiss National Science Foundation (to C.D.). The Center for Structural studies is funded by the DFG (Grant number 417919780 and INST 208/761-1 FUGG to S.S.).

## Notes

### Competing Interest Statement

The authors have declared no competing interest.

